# Total RNA-sequencing reveals multi-level microbial community changes and functional responses to wood ash application in agricultural and forest soil

**DOI:** 10.1101/621557

**Authors:** Toke Bang-Andreasen, Muhammad Zohaib Anwar, Anders Lanzén, Rasmus Kjøller, Regin Rønn, Flemming Ekelund, Carsten Suhr Jacobsen

**Affiliations:** Department of Environmental Science, Aarhus University, RISØ campus, Roskilde, Denmark; Department of Biology, University of Copenhagen, Copenhagen, Denmark; Department of Conservation of Natural Resources, NEIKER-Tecnalia, Bizkaia Technology Park, Derio, Spain; AZTI-Tecnalia, Herrera Kaia, Pasaia, Spain; IKERBASQUE, Basque Foundation for Science, Bilbao, Spain; Key Laboratory of Urban Environment and Health, Institute of Urban Environment, Chinese Academy of Sciences, Xiamen, China; Arctic Station, University of Copenhagen, Qeqertarsuaq, Greenland

**Author notes:** Corresponding Author: Flemming Ekelund.

**Keywords:** Metatranscriptomics, Total RNA, wood ash, soil micro-biome, biodiversity, soil biota, mRNA, rRNA, renewable energy, protozoa

## Abstract

Recycling of wood ash from energy production may counteract soil acidification and return essential nutrients to soils. However, wood ash amendment affects soil physicochemical parameters that control composition and functional expression of the soil microbial community. Here, we applied Total RNA-sequencing to simultaneously assess the impact of wood ash amendment on the active soil microbial communities and the expression of functional genes from all microbial taxa. Wood ash significantly affected the taxonomic (rRNA) as well as functional (mRNA) profiles of both agricultural and forest soil. Increase in pH, electrical conductivity, dissolved organic carbon and phosphate were the most important physicochemical drivers for the observed changes. Wood ash amendment increased the relative abundance of the copiotrophic groups Chitinonophagaceae (Bacteroidetes) and Rhizobiales (Alphaproteobacteria) and resulted in higher expression of genes involved in metabolism and cell growth. Finally, Total RNA-sequencing allowed us to show that some groups of bacterial feeding protozoa increased concomitantly to the enhanced bacterial growth, which shows their pivotal role in the regulation of bacterial abundance in soil.

## 1. Introduction

Wood ash from energy production is often considered a waste product (Vance, 1996; Demeyer *et al.*, 2001) despite that recycling of wood ash may have beneficial effects as it counteracts acidification and returns essential nutrients to soil (Demeyer *et al.*, 2001; Augusto *et al.*, 2008). Wood combustion is becoming more popular in several countries and increased reuse of wood ash as soil amendment holds the potential to improve the sustainability of this practice (Karltun *et al.*, 2008; Huotari *et al.*, 2015). However, wood ash application affects several soil physicochemical parameters important to the structure and function of microbial communities; e.g. pH, electrical conductivity and dissolved organic carbon (DOC), (Ohno & Susan Erich, 1990; Demeyer *et al.*, 2001; Pitman, 2006; Augusto *et al.*, 2008; Maresca *et al.*, 2017; Hansen *et al.*, 2017). As the soil micro-biota carries out an array of key biochemical processes (Blagodatskaya & Kuzyakov, 2013), knowledge of its response to disturbance is important; not least in production soils due to potential impact on soil fertility.

The soil micro-biome, which includes prokaryotes as well as micro-eukaryotes, is one of the most diverse and complex biomes on Earth. It has a pivotal role in nutrient cycling and carbon sequestration and is a key component in the maintenance of soil fertility of managed ecosystems (Wall *et al.*, 2012; Fierer, 2017). Wood ash amendment causes changes in soil micro-biome composition, activity and quantity (Perkiömäki & Fritze, 2002; Aronsson & Ekelund, 2004; Huotari *et al.*, 2015). Ash amendment induces changes in community structure followed by increased microbial activity and growth, which is usually explained by the increased soil pH brought about by the alkaline oxides in the ash (Cruz-Paredes *et al.*, 2017; Vestergård *et al.*, 2018). Still, some studies show no or only minor microbial response to wood ash application (Aronsson et al. 2004; Huotari et al. 2015). Only few studies have concomitantly analysed microorganisms from all domains of life (i.e. Archaea, Bacteria, Eukaryotes) and most of these rely on cultivation, model organisms or molecular fingerprinting, which only provide limited resolution of taxonomical and functional responses. Total RNA-sequencing, or metatranscriptomics, makes it possible to investigate active soil microbial communities from all domains of life, incl. their transcriptional activity, simultaneously. Because total RNA-sequencing allows for the study of immediate regulatory responses to environmental changes (Carvalhais *et al.*, 2012), it has also proven useful in the assessment of active microbial communities’ functional roles in soil (Urich *et al.*, 2008; Hultman *et al.*, 2015; Epelde *et al.*, 2015; Geisen *et al.*, 2015; Schostag *et al*., 2019).

We therefore aimed to investigate how the active soil prokaryotic and micro-eukaryotic communities in agricultural and forest soil responded structurally and functionally (transcriptional) to wood ash application. Both soil types are relevant for large scale application of wood ash. We applied wood ash in concentrations corresponding to field application of 0, 3, 12 and 90 t ha^−1^, where 3 t ha^−1^ is the currently allowed dose in Scandinavian countries. We expected wood ash to increase soil pH, electrical conductivity and dissolved organic carbon and therefore hypothesised that (I) the pH increase would favour bacteria more than fungi, (II) the nutrients in the wood ash would benefit the copiotrophic microbial groups, (III) multitrophic responses would appear gradually over time after wood ash application, and (IV) that microbial stress responses would be observable in the transcriptome.

## 2. Materials & Methods

### 2.1. Soils and wood ash

We used two contrasting soils for the experiment. The first was a loamy sand (Typic Hapludult) from the plough layer (0-10 cm) of an agricultural field (Research Center Foulum, DK, 56°29’42”N 9°33’36”E). The other was from the O-horizon (0-10 cm) of a forest (Gedhus, DK, 56°16’38”N 09°05’12”E). The forest is a second-generation Norway spruce stand (*Picea abies* (L.) Karst.) on Podzol heathland. Qin *et al.* (2017) provide soil characteristics for both soils. On both sites, we removed visible plant parts before taking ten 100 g soil samples within a 30 m^2^ area. The 10 samples from each site were sieved (4 mm), pooled and stored in the dark at 4 °C until further processing.

Wood ash was a mixture of bottom-and fly-ash from a heating plant (Brande, Denmark) produced by combustion of wood chips from predominantly coniferous trees. We homogenized the ash by sieving (2 mm). Maresca et al. (2017) provide a list of mineral nutrients and heavy metals in the ash.

### 2.2. Microcosm set-up and incubation

We prepared microcosms in triplicates of 50 g soil in 250 ml sterilised airtight glass jars. We mixed the ash thoroughly with soil to ash concentrations corresponding to field application of 0, 3, 12 and 90 t ash ha^−1^. The water content was adjusted to 50 % of the water holding capacity of the two soils. We prepared 12 microcosms for each soil-ash combination to allow four destructive samplings; i.e. a total of 96 microcosms. Samples were also collected at the start of the experiment. Microcosms were incubated at 10 °C in the dark and all microcosms were opened once a week inside a LAF-bench to maintain aerobic conditions.

### 2.3. Physicochemical soil parameters

At destructive sampling, after 3, 10, 30 and 100 days of incubation, we prepared soil extracts from 15 g soil and 75 ml sterile ddH_2_O followed by 1 h shaking and settling for 0.5 h. In the supernatant, we measured electrical conductivity using a TetraCon 325 electrode adapted to a conductivity meter Cond 340i (WTW, Weilheim, Germany) and pH using a pH electrode (Sentix Mic) connected to pH meter Multi 9310 (WTW). The remaining supernatant was filtered (5C filters, Advantec, Tokyo, Japan; 1 μm pore size) and analysed for dissolved organic carbon (DOC), nitrate (NO_3_^−^), ammonium (NH_4_^+^) and phosphate (PO_4_^3-^). DOC concentrations were determined on a TOC-5000A (Shimadzu, Kyoto, Japan). Nitrate, ammonium and phosphate concentrations were determined by flow injection analysis (FIAstar™ 5000, FOSS, Hillerød, Denmark) following manufacturer’s instructions.

### 2.4. Nucleic acid extraction, qPCR and library preparation for sequencing

RNA and DNA were co-extracted from 2 g soil samples using the RNA PowerSoil Total RNA Isolation Kit (MOBIO, Carlsbad, USA) in combination with DNA Elution Accessory Kit (MOBIO) following manufacturer’s protocol. Agricultural soil amended with the highest ash concentration had an RNA yield below detection limit and was not sequenced.

We quantified 16S rRNA and ITS2 gene copies (DNA level) using qPCR. 16S rRNA genes were amplified in technical duplicates using a CFX Connect (Bio-Rad, Richmond, USA). We used a dilution series of genomic DNA from *Escherichia coli* K-12 (with 7 copies of 16S rRNA genes) as a standard (Blattner *et al.*, 1997). The master-mix consisted of 2 µl bovine serum albumin (BSA) (20mg/ml; BIORON, Ludwigshafen, Germany), 10 µl SsoFast EvaGreen Supermix (Bio-Rad), 0.8 μl of primer 341f (5′-CCTAYGGGRBGCASCAG-3′), 0.8 μl of primer 806r (5′-GGACTACNNGGGTATCTAAT-3′) (Hansen *et al.*, 2012); 1 μl of 10× diluted template, and 5.4 µl sterile DEPC-treated water. PCR conditions for 16S rRNA gene amplification were 98°C for 15 min, followed by 35 cycles of 98°C for 30 s, 56°C for 30 s, and 72°C for 30 s (with fluorescence measurements) and ending with 72°C for 7 min and production of melt curves. The PCR efficiencies for the 16S assays were 96.1±1.0% (SEM, n=3) with R^2^ = 0.99±0.001. ITS gene copies were quantified as described for the 16S rRNA above with minor modifications: Vector cloned ITS2 DNA regions from *Aureobasidium pullulans* were included as standards, primers used were gITS7 (5′-GTGARTCATCGARTCTTTG-3′ (Ihrmark *et al.*, 2012)) and ITS4 (5′-TCCTCCGCTTATTGATATGC-3′ (White *et al.*, 1990)), annealing temperature was 60 °C and 40 amplification cycles were used. The PCR efficiencies for the ITS assays were 106.0±4.6% with R^2^ = 0.99±0.003.

Prior to Total RNA library building, we removed potential DNA carryovers using the DNase Max Kit (MOBIO) following manufacturer’s protocol. Successful DNA removal of RNA extracts were tested with the 16S qPCR protocol described above but with 50 amplification cycles: All DNase treated RNA extracts had higher or equal Cq values than the negative samples (sterile DEPC-treated water as template) and DNA was thereby not present.

Quality of the DNase treated RNA was tested using RNA 6000 Nano Kit (Agilent, Santa Clara, USA) on a 2100 Bioanalyzer System (Agilent) following manufacturer’s protocol (Average RIN number was 7.85±0.13 (SEM, n=69)

Subsequently, DNase treated RNA extracts from time points 0, 3, 30 and 100 days were fragmented into ∼150 bp segments and prepared for sequencing using the NEBNext Ultra Directional RNA Library Prep Kit for Illumina in combination with the NEBNext Multiplex Oligos for Illumina (New England BioLabs, Ipswich, USA) according to the manufacturer’s protocol. We sequenced the resulting metatranscriptome libraries using HiSeq 2500 (Illumina Inc., San Diego, USA) in high output mode (8 HiSeq lanes, 125bp, paired end reads) at the National High-throughput DNA Sequencing Centre (Copenhagen, Denmark).

### 2.5. Bioinformatic processing

We obtained a total of 3.3 billion paired sequences (SRA accession number: PRJNA512608) and processed them through the following bioinformatic pipeline. Adapters, poly-A tails, sequences shorter than 60 nt and nucleotides with phred score below 20 at the 5’ and 3’ end of sequences were removed using Cutadapt v.1.9.1 (Martin, 2011). Five samples were removed prior to subsequent processing due to low quality of reads (one replicate of 3 t ha^−1^, day 100 from the agricultural soil and two replicates of 0 t ha^−1^, day 0 and two replicates 0 t ha^−1^, day 100 from the forest soil). Sequences were then sorted into small subunit (SSU) rRNA, large subunit (LSU) rRNA and non-rRNA sequences using SortMeRNA v.2.1 (Kopylova *et al.*, 2012).

#### 2.5.1. rRNA

A subset of 1.5 million randomly chosen SSU rRNA sequences per sample were assembled into longer SSU rRNA contigs using EMIRGE (Miller *et al.*, 2011). Contigs were taxonomically classified using CREST (Lanzén *et al.*, 2012) and rRNA reads were mapped to resulting EMIRGE contigs using BWA (Li & Durbin, 2009), as in Epelde *et al.* (2015), resulting in a table of taxonomically annotated read abundance across samples (Supplementary Datasheet 1).

#### 2.5.2. mRNA

A combined pool of non-ribosomal sequences from all samples was assembled using trinity v.2.0.6 (Grabherr *et al.*, 2011). From the resulting assembled contigs, non-coding RNA contigs were filtered away by aligning contigs to the Rfam database v.12.0 (Nawrocki *et al.*, 2015) using cmsearch v.1.1.1 with a significant e-value threshold of <10^−3^. Input sequences used for non-ribosomal RNA assembly were then mapped to coding mRNA contigs. We normalized the contigs by removing those with relative expression lower than 1 out of the number of sequences in the dataset with least number of sequences. EMBOSS (Rice *et al.*, 2000) was used to search six possible open reading frames (ORFs) of the contigs. SWORD (Vaser *et al.*, 2016) was used to align ORFs against the Md5nr protein database (Wilke *et al.*, 2012). The output was then parsed with custom Python scripts and filtered hits with minimum e-value of 10^−5^ as threshold. Best hit for each contig was then selected based on alignment statistics and annotated against the eggnog hierarchical database v.4.5 (Jensen *et al.*, 2008). The output was an abundance table of numbers of sequences assigned to groups of different functional genes (COGs) (Supplementary Datasheet 2).

### 2.6. Statistical analysis and data processing

Statistical validation for both taxonomy and functional abundance was done in R v.3.4.0 (R Core Team, 2015) using *vegan* (Oksanen *et al.*, 2008). The rRNA abundance was converted into relative abundance and collapsed taxonomically into Archaea, Bacteria and Eukaryota. We further grouped Eukaryota into Fungi, Metazoa, and protists (with main focus on bacterivorous protozoa). We calculated Richness and Shannon diversity on the total number of rRNA contigs and abundance of sequence reads mapped to them. Non-metric multidimensional scaling (NMDS) was carried out using Bray-Curtis dissimilarities of community composition between samples. Soil physicochemical parameters were fitted to the resulting NMDS using the function *envfit*. Variables explaining overall differences in community composition were evaluated using the function *Adonis*, which performs permutational analysis of variance (PERMANOVA; 10,000 permutations) using Bray-Curtis dissimilarities as response variable. A forward selection strategy was carried out to only include explanatory variables with significant p-values in *Adonis* models.

Significant effects of wood ash amendment and incubation time on taxonomic groups were determined using non-parametric Kruskal-Wallis tests (due to the non-normal distribution of taxon abundances). To separate the pronounced changes in community responses observed at the 90 t ha^−1^ amendment in the forest soil from the less pronounced changes observed at 0-12 t ha^−1^, we performed Kruskal-Wallis tests with wood ash concentration as independent variable for both the range of 0-12 t ha^−1^ and 0-90 t ha^−1^. We also used Kruskal-Wallis to test the effect of time on differential abundances within the wood ash concentrations separately. P-values were adjusted for false discovery rate (FDR) using the Benjamini–Hochberg method in all tests. NMDS on Bray Curtis dissimilarities of gene compositions and *Adonis* testing were carried out as described above. Gene counts between samples were normalized using the DESeq2 algorithm (Love *et al.*, 2014). Significantly differentially expressed genes were analysed using the DESeq2 module of SARTools (Varet *et al.*, 2016). These analyses were conducted by pairwise comparisons of gene transcription levels between samples of increasing wood ash concentration to control samples (0t ha^−1^) at different incubation times. For the forest samples at time 100 days, only one replicate remained for the 0 t ha^−1^ treatment. Therefore we compared instead the 12 and 90 t ha^−1^ to the 3 t ha^−1^.

We used linear Pearson regression to test for significant correlations between wood ash concentration and time against measured physicochemical parameters. Additionally, we performed two-way ANOVAs with Tukey’s post-hoc tests using wood ash concentration and time as explanatory variables, with all physicochemical parameters as dependent variables. Variance homogeneity was tested using Levenes’s test and normal distribution of data was tested using the Shapiro-Wilk test in combination with QQ-plots prior to ANOVA tests.

We used a significance level of 0.05, unless otherwise explicitly mentioned, and the results section provide descriptions at this significance level.

## 3. Results

### 3.1. Physicochemical parameters

Soil pH, electrical conductivity and DOC correlated positively with wood ash concentration for both soils (Table 1). For the 90 t ha^−1^ ash amendment, soil pH increased from 6.4 to 11.5 in the agricultural and from 4.1 to 8.5 in the forest soil (Supplementary Figure 1). Similarly, the 90 t ha^−1^ resulted in 15 and 19-fold increases in electrical conductivity for the agricultural and forest soil, respectively. In the agricultural soil, ammonium increased with time in samples both with and without ash amendment, while nitrate showed no significant changes. In the forest soil, ammonium and nitrate increased after 3 days in the 90 t ha^−1^ amendment, followed by a decrease after 30 days. In the other treatments, increased concentrations were observed during the entire incubation period. In both soils, concentrations of dissolved phosphate increased up to 12 t ash ha^−1^ followed by a decrease at 90 t ha^−1^.

**Table 1:**
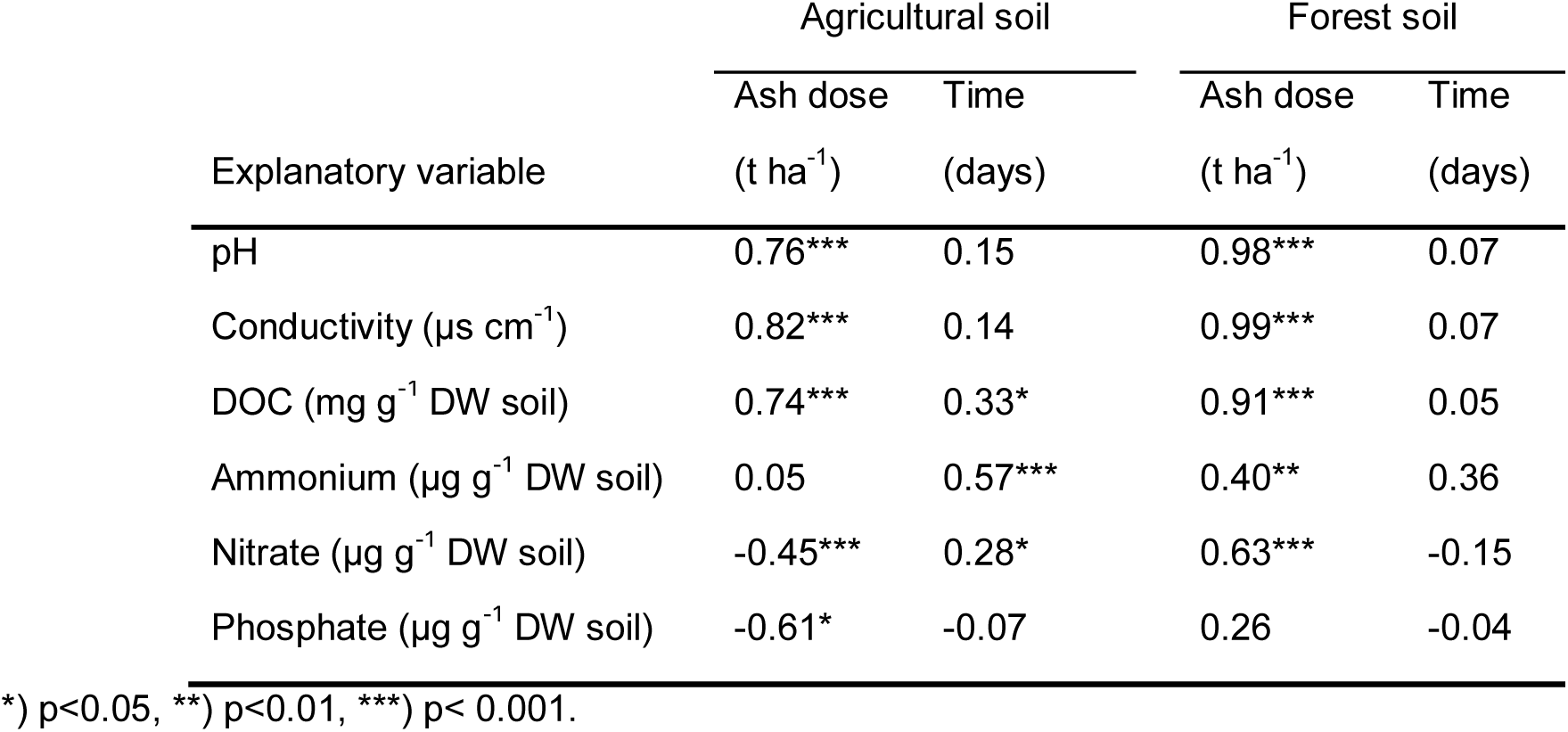
Pearson correlation values (r) and associated significance levels between ash dose (field equivalents 0, 3, 12 and 90 t ha-1) and incubation time, and soil physicochemical parameters.

| Explanatory variable | Agricultural soil |  | Forest soil |  |
| --- | --- | --- | --- | --- |
|  | Ash dose<br>(t ha<sup>-1</sup>) | Time<br>(days) | Ash dose<br>(t ha<sup>-1</sup>) | Time<br>(days) |
| pH | 0.76*** | 0.15 | 0.98*** | 0.07 |
| Conductivity (µs cm<sup>-1</sup>) | 0.82*** | 0.14 | 0.99*** | 0.07 |
| DOC (mg g<sup>-1</sup> DW soil) | 0.74*** | 0.33* | 0.91*** | 0.05 |
| Ammonium (µg g<sup>-1</sup> DW soil) | 0.05 | 0.57*** | 0.40** | 0.36 |
| Nitrate (µg g<sup>-1</sup> DW soil) | -0.45*** | 0.28* | 0.63*** | -0.15 |
| Phosphate (µg g<sup>-1</sup> DW soil) | -0.61* | -0.07 | 0.26 | -0.04 |

\*) p<0.05, \*\*) p<0.01, \*\*\*) p< 0.001.

### 3.2. Quantitative PCR

Prokaryotic abundance (number of 16S rRNA gene copies) increased in the agricultural soil after the wood ash application of 12 t ha^−1^, but decreased after application of 90 t ha^−1^ (Figure 1). Fungal abundance (number of ITS copies) remained fairly unchanged over time regardless of ash application with the exception of an increase after 100 days at 90 t ha^−1^. In the forest soil prokaryotic abundance increased over time for all treatments (Figure 1); however, addition of 12 and 90 t ha^−1^ resulted in a stronger increase. The fungal abundance in the forest soil showed higher abundance for most of the period with wood ash concentrations of 90 t ha^−1^.

**Figure 1:**
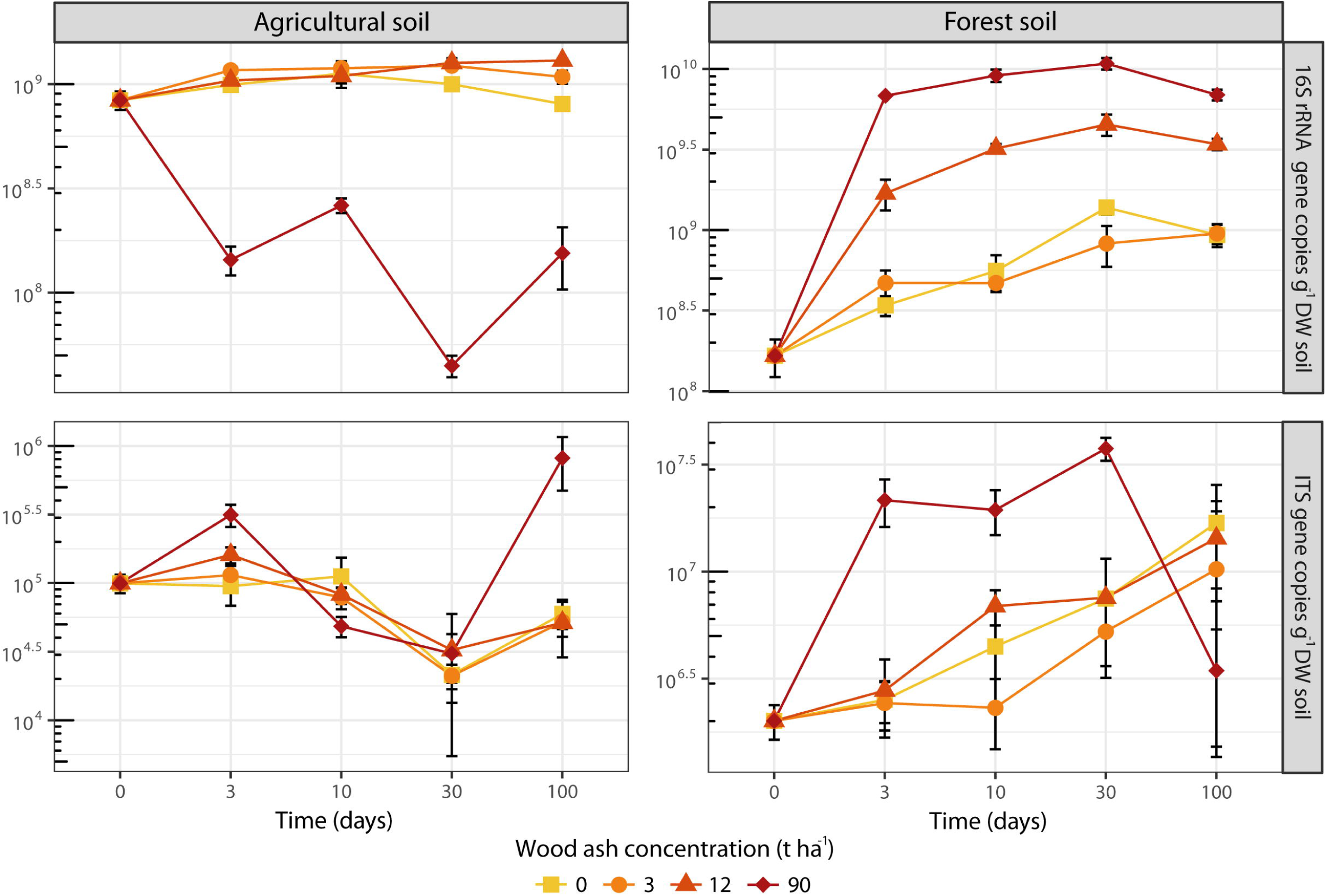
Numbers of 16S rRNA gene copies (top row) and ITS gene copies (bottom row) per g-1 DW of the agricultural soil (left panel) and the forest soil (right panel) across wood ash concentrations and incubation times. Symbols represents averages with SEM (n=3). The presented data are results from qPCR on DNA. Note logarithmic y-axes and different ranges of values on y-axes.

### 3.3. rRNA - Community composition

The number of unique rRNA contigs ranged from 1,216 – 5,931 per sample and originated from all domains of life. Community composition differed significantly (p < 0.001; R^2^ = 0.86; *Adonis*) between the two soil types. For forest soil, amendment with 90 t ha^−1^ resulted in highly altered community composition (Figure 2) compared to 0-12 t ha^−1^. Though less pronounced, changes from 0-3-12 t ha^−1^ were also clearly visible for both soil types (Figure 2). Moreover, microcosms for particular soil type/ash dose combinations were clearly separated by sampling times (Figure 2).

**Figure 2:**
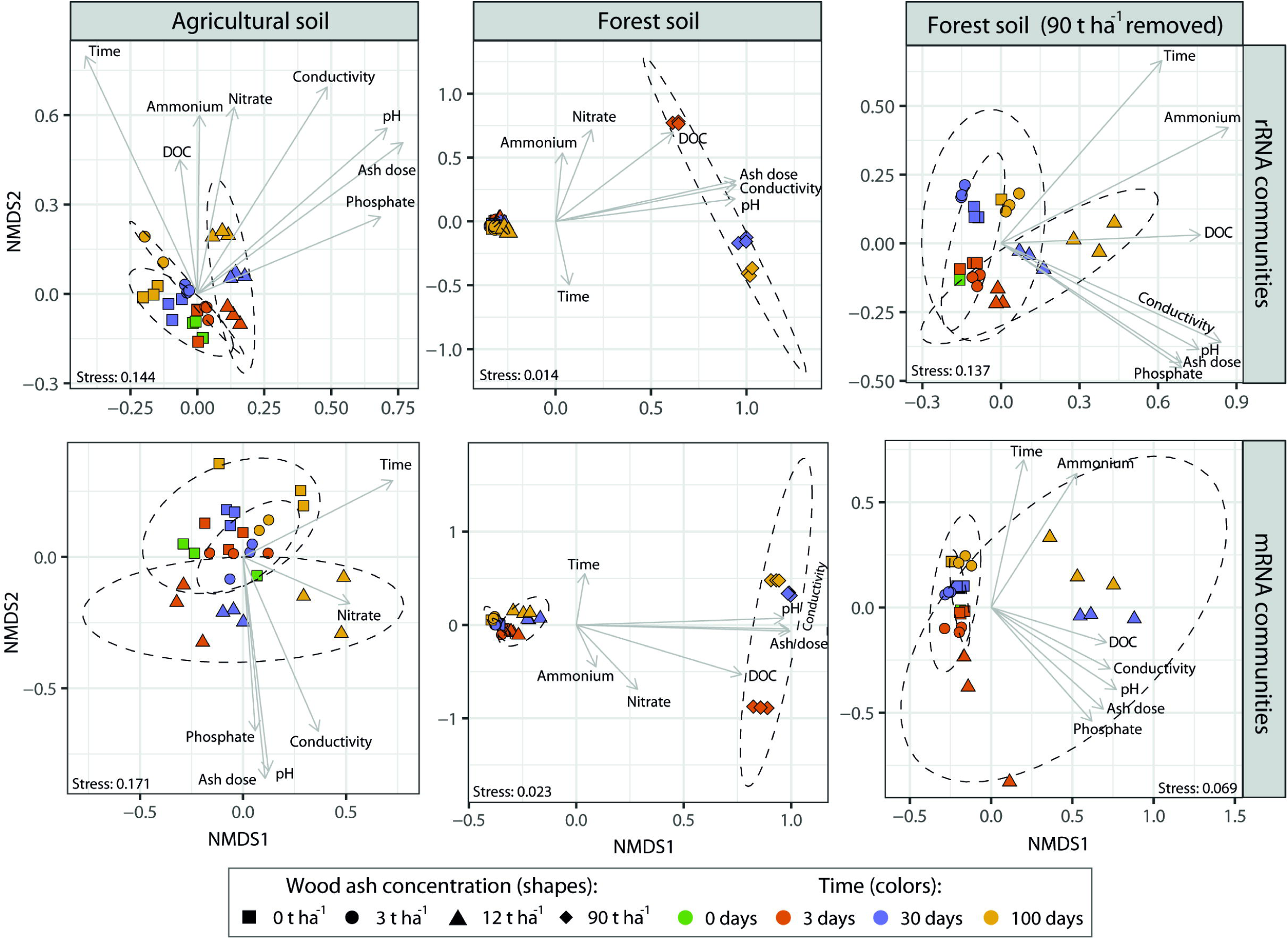
NMDS plots based on Bray-Curtis dissimilarities of the taxonomic (rRNA; top row) and functional (mRNA; bottom row) profiles of an agricultural soil and a forest soil amended with wood-ash. Dashed lines represent 95% confidence ellipses around samples with same wood ash concentration. Arrows indicate the direction of fitted physiochemical parameters (using envfit function; only significant parameters shown) onto the NMDS ordination space (longer arrows indicate better fit). To improve the resolution of the forest soil at wood ash concentrations 0-12 t ha-1, we removed the 90 t ha-1 samples and repeated the analysis (rightmost two panels).

In both soils, wood ash dose, incubation time, pH and electrical conductivity correlated to the transformed NMDS community space (Figure 2). Optimized *Adonis* models (Table 2) supported that wood ash concentration, time, pH, and electrical conductivity together significantly explained the variation in microbial communities after ash application in both soils. Additionally, dissolved phosphate significantly explained the variation in microbial communities in both soils up to 12 t ha^−1^ ash amendments and DOC, ammonium and nitrate in the forest soil.

**Table 2:**
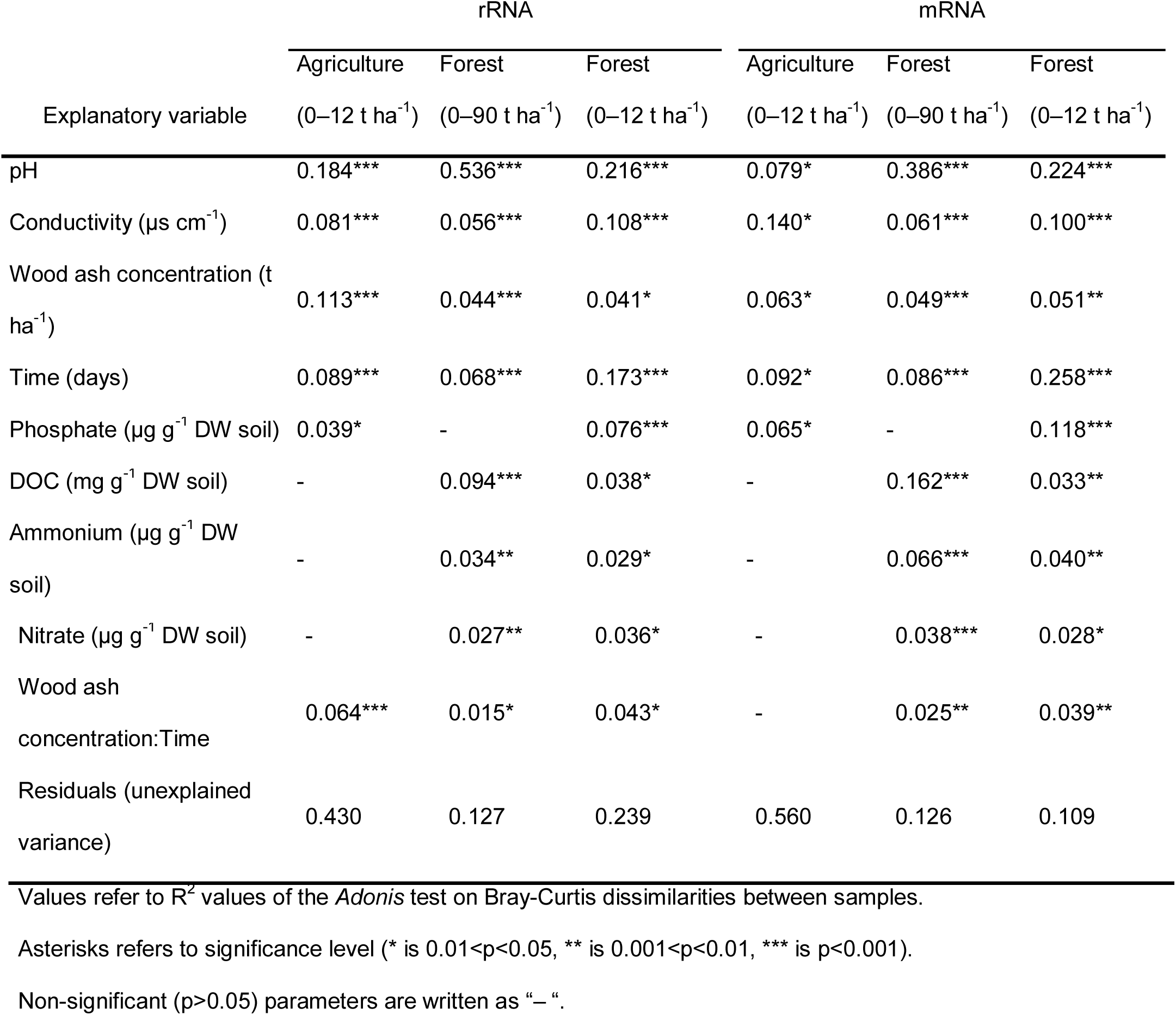
Explanatory strength of physiochemical variables on rRNA and mRNA dissimilarity profiles of the two soils after ash amendment tested using Permutational Multivariate Analysis Of Variance (*Adonis*)

| Explanatory variable | rRNA |  |  | mRNA |  |  |
| --- | --- | --- | --- | --- | --- | --- |
|  | Agriculture<br>(0–12 t ha<sup>-1</sup>) | Forest<br>(0–90 t ha<sup>-1</sup>) | Forest<br>(0–12 t ha<sup>-1</sup>) | Agriculture<br>(0–12 t ha<sup>-1</sup>) | Forest<br>(0–90 t ha<sup>-1</sup>) | Forest<br>(0–12 t ha<sup>-1</sup>) |
| pH | 0.184*** | 0.536*** | 0.216*** | 0.079* | 0.386*** | 0.224*** |
| Conductivity (µs cm<sup>-1</sup>) | 0.081*** | 0.056*** | 0.108*** | 0.140* | 0.061*** | 0.100*** |
| Wood ash concentration (t ha<sup>-1</sup>) | 0.113*** | 0.044*** | 0.041* | 0.063* | 0.049*** | 0.051** |
| Time (days) | 0.089*** | 0.068*** | 0.173*** | 0.092* | 0.086*** | 0.258*** |
| Phosphate (µg g<sup>-1</sup> DW soil) | 0.039* | - | 0.076*** | 0.065* | - | 0.118*** |
| DOC (mg g<sup>-1</sup> DW soil) | - | 0.094*** | 0.038* | - | 0.162*** | 0.033** |
| Ammonium (µg g<sup>-1</sup> DW soil) | - | 0.034** | 0.029* | - | 0.066*** | 0.040** |
| Nitrate ($\mu\text{g g}^{-1}$ DW soil) | - | 0.027** | 0.036* | - | 0.038*** | 0.028* |
| Wood ash<br>concentration:Time | 0.064*** | 0.015* | 0.043* | - | 0.025** | 0.039** |
| Residuals (unexplained<br>variance) | 0.430 | 0.127 | 0.239 | 0.560 | 0.126 | 0.109 |

Values refer to $R^2$ values of the *Adonis* test on Bray-Curtis dissimilarities between samples.
Asterisks refers to significance level (\* is $0.01 < p < 0.05$ , \*\* is $0.001 < p < 0.01$ , \*\*\* is $p < 0.001$ ).
Non-significant ( $p > 0.05$ ) parameters are written as “-”.

### 3.4. rRNA – Taxonomic distribution and diversity

A majority (85%) of sequence reads, mapped to contigs, could be annotated to order rank (99% to phylum and 97% to class rank) (Figure 3A). Fewer sequences could be assigned lower taxonomic ranks (60% and 27% to family and genus level, respectively). Therefore, we evaluated possible significant differences in abundance of taxa at order level (see Supplementary Datasheet 3 and 4 for p-values and averages of relative abundances, respectively). Richness and Shannon diversity decreased in the unamended agricultural soil over time, while ash amendments of 3 and 12 t ha^−1^ counteracted this decrease (Figure 3B). In the forest soil these measures generally remained unchanged up to 12 t ha^−^ 1 amendments (with a single exception of increased richness at 3 t ha^−1^ after 100 days of incubation), while the 90 t ha^−1^ amendment caused reduction of Shannon diversity.

**Figure 3:**
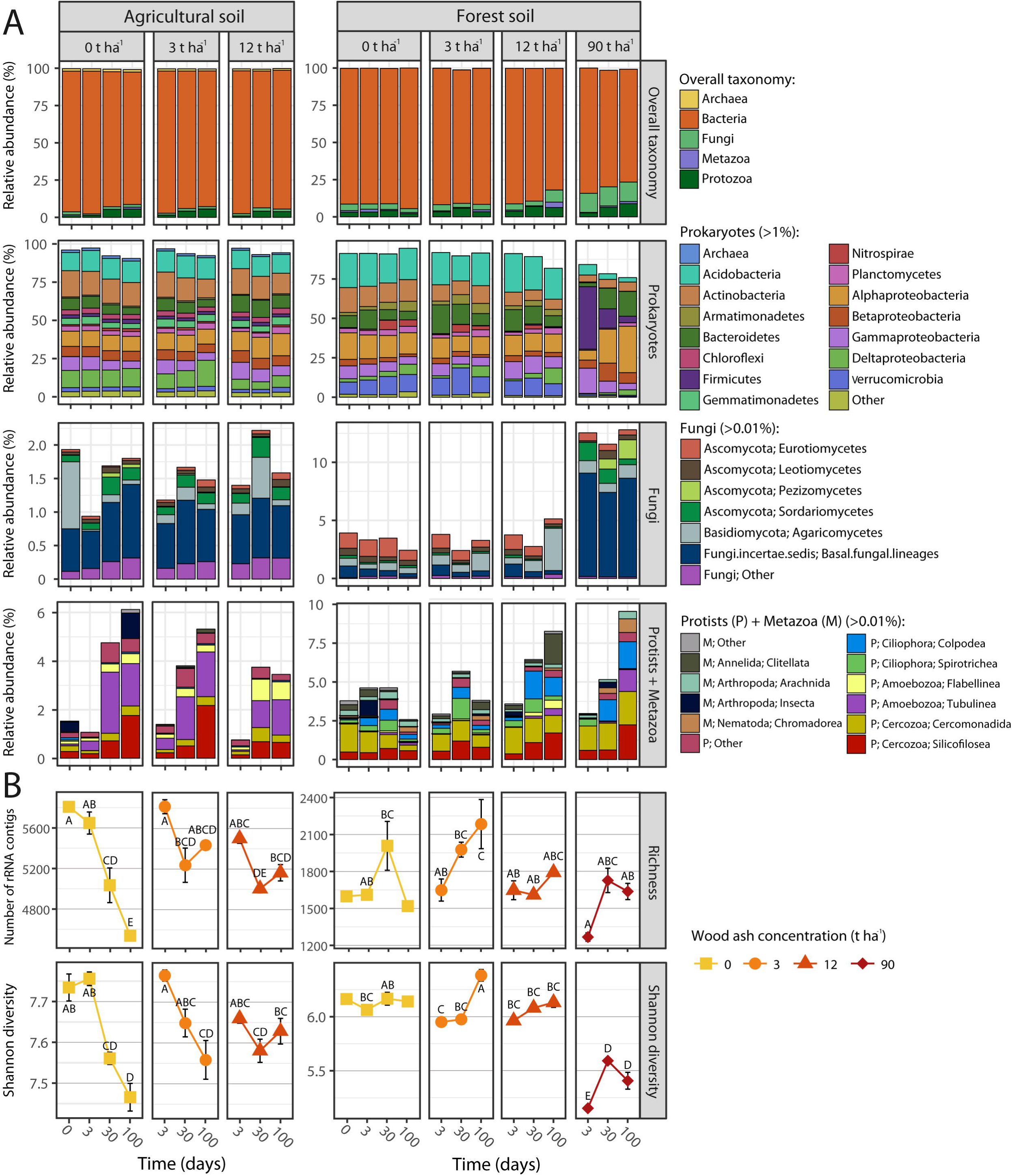
Community composition and diversity across the two soils at increasing wood ash amendment and incubation times based on PCR-free, total RNA-Seq. (A). The most abundant taxonomic groups (cutoff levels of average relative abundances are shown in legend header) are presented in upper panel (Overall taxonomy); i.e. Archaea, Bacteria, Fungi, Protists, and Metazoa. Bars represente averages of triplicates (excluding agricultural soil 3 t ha^−1^ at 100 days (n=2), forest soil 0t ha^−1^ at 0 days (n=1) and forest soil 0t ha^−1^ at 100 days (n=1)). (B) Richness and Shannon diversity. Statistically significant different richness and diversity measures (p < 0.05) between samples within each measure and soil is indicated by different letters. Symbols represent averages, as described for the bar plots.

#### 3.4.1. Prokaryotic community

In both soil types, the relative abundance of Chitinophagaceae (Bacteroidetes) increased with wood ash application (Figure 3A). In the agricultural soil, ash-amendment also caused increases in Alphaproteobacteria and Betaproteobacteria. In the forest soil, the 3 and 12 t ha^−1^ ash-amendments increased Myxococcales (Deltaproteobacteria), while Acidimicrobiia (Actinobacteria) decreased.

In the forest soil, the 90 t ha^−1^ ash-amendment resulted in major prokaryotic community changes. Actinobacteria, Acidobacteria, Armatimonadetes, and Verrucomicrobia decreased, while Firmicutes, Bacteroidetes and Proteobacteria increased. Firmicutes dominated after 3 days, with *Paenibacillus* as most abundant with relative abundance of 21.3%, followed by a gradual decrease towards 1.1% after 100 days. Similarly, Gammaproteobacteria decreased during incubation after an initial increase. Chitinophagaceae and Rhizobiales (Alphaproteobacteria) showed the opposite temporal trend and were most abundant after100 days.

#### 3.4.2. Fungal community

The 3 and 12 t ha^−1^ ash amendments did not affect fungal community composition in the agricultural soil (Figure 3A). In the forest soil no major changes were found at low amendments, while application of 90 t ha^−1^ resulted in increase in *Mortierella*, Hypocreales (Sordariomycetes) and *Peziza* (Pezizomycetes).

#### 3.4.3. Micro-eukaryotic community

In the agricultural soil, the relative abundances of Tubulinea (Amoebozoa), Thaumatomonadida (Cercozoa) and Silicofilosea (Cercozoa) increased over time in all treatments (Figure 3A). In the forest soil, *Colpoda* (Ciliophora) increased with time in all treatments, though more pronounced at higher wood ash amendments. Further, Tubulinea (Amoebozoa), Heteromitidae (Cercozoa) and Silicofilosea (Cercozoa) increased in the 12 and 90 t ha^−1^ amendments.

### 3.5. mRNA - Functional genes

A total of 0.9 million sequences were mapped to 463 mRNA contigs (Supplementary Figure 2 and Supplementary Datasheet 2). The two soils possessed distinct pools of expressed genes (p < 0.001; R^2^ = 0.82; A*donis*). Overall, Bray-Curtis dissimilarities, based on mRNA profiles, and fitting of physicochemical parameters to these, revealed similar trends as for rRNA taxonomic communities (Figure 2 and Table 2).

In the agricultural soil, we observed only minor functional gene responses to time and ash-amendment, while more genes were differentially expressed in the forest soil, especially at the 90 t ha^−1^ amendment (Figure 4; full list of differential expressed genes in Supplementary Datasheet 5). Of the well characterized genes, four functional categories contained most of the differentially expressed genes; i.e. “Post-translation modification, protein turnover, and chaperones”, “Transcription”, “Replication, recombination and repair” and “Carbohydrate transport and metabolism”. Furthermore, genes related to stress responses increased mainly in the forest soil at 90 t ha^−1^ ash amendments (Supplementary Figure 3).

**Figure 4:**
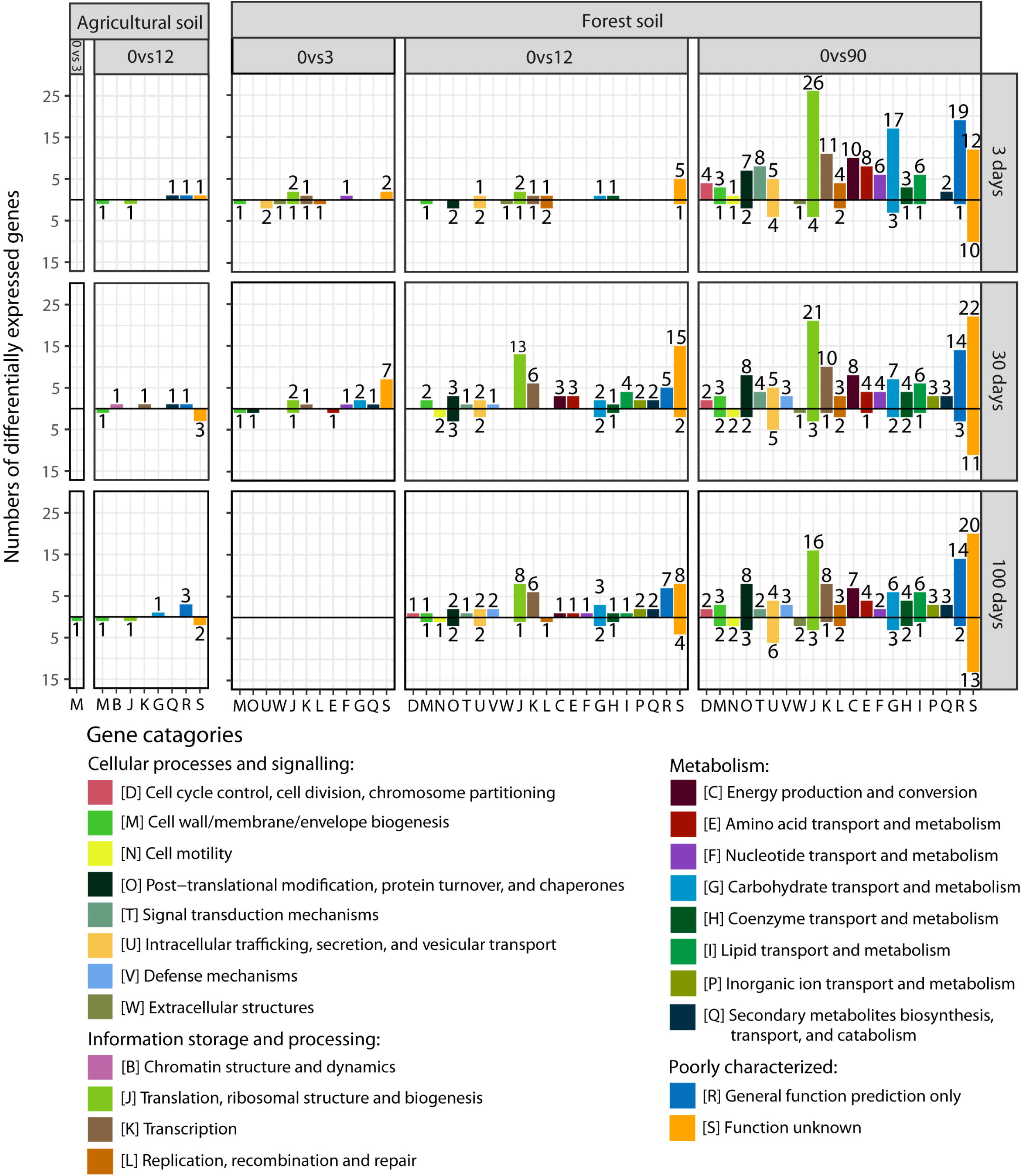
Numbers of differentially expressed genes within functional categories across agricultural and forest soil by pairwise comparisons of gene transcription levels between samples of increasing wood ash concentration to reference samples without ash-amendment at different incubation times. “0vs3, “0vs12” and “0vs90” denote the wood ash doses compared, i.e. wood ash dose 0 t ha-1 compared to 3 t ha-1 is written “0vs3”. Increasing and decreasing gene transcription levels are presented above and below the black horizontal zero-line, respectively. The pairwise comparisons for forest soil, 100 days, were carried out using 3 t ha-1, 100 days, as reference samples because only one replicate was acquired from the 0 t ha-1, 100 days, samples (hence the empty plot in 0vs3, 100 days, forest plot). Digits above/below bars represent the number of differentially expressed genes within a gene category.

## 4. Discussion

Here, we present the first detailed analysis of changes in soil microbial prokaryotic and eukaryotic communities after amendment with ash using the Total RNA-sequencing procedure.

### 4.1. Bacterial responses to wood ash application

The general copiotrophic groups of bacteria, i.e. Bacteroidetes, Alphaproteobacteria and Betaproteobacteria were stimulated by wood ash application. Members of Bacteroidetes benefit from wood ash application (Noyce *et al.*, 2016; Bang-Andreasen *et al.*, 2017); they are initial metabolizers of labile carbon and respond positively to increased soil pH and electrical conductivity (Fierer *et al.*, 2007; Lauber *et al.*, 2009; Kim *et al.*, 2016). Alpha- and Betaproteobacteria are also generally copiotrophic (Fierer *et al.*, 2007; Cleveland *et al.*, 2007); Betaproteobacteria thrive in soils with higher pH (Kim *et al.*, 2016), whereas Alphaproteobacteria are favoured at high N availability (Nemergut *et al.*, 2010; Fierer *et al.*, 2012).

Acidobacteria and Verrucomicrobia declined after the 90 t ha^−1^ amendment to the forest soil. These phyla are considered oligotrophic (Fierer *et al.*, 2007; Bergmann *et al.*, 2011; Ramirez *et al.*, 2012; Cederlund *et al.*, 2014; Kielak *et al.*, 2016) and Acidobacteria are generally most abundant under acidic conditions (Rousk *et al.*, 2010; Kielak *et al.*, 2016). Thus, increases in pH, bioavailable DOC and nutrients induced by wood-ash allow copiotrophic groups to thrive on the expense of oligotrophic groups. The shift towards a more copiotrophic dominated community after ash amendment was further supported by the mRNA profile of the soil. Here, an increasing number of functional genes involved in metabolism and cell growth (“Translation”, “Transcription” and “Replication”) showed significant higher transcription levels.

Of the Bacteroidetes, Chitinonophagaceae showed the strongest positive response to wood ash application. Members of this family can degrade a broad spectrum of carbon compounds (Kämpfer *et al.*, 2006; Hanada *et al.*, 2014). Thus, they are well suited for the ash-induced increased DOC availability. Rhizobiales dominated the increasing Alphaproteobacterial fraction of the forest soil after ash amendment. They are copiotrophs (Starke *et al.*, 2016; Lladó & Baldrian, 2017) and can degrade organic pollutants and cope with heavy metals (Teng *et al.*, 2015). Probably advantageous properties, as the wood ash induces increase of heavy metals and nutrients in the soils. Deltaproteobacterial Myxococcales responded positively to wood ash amendment in the forest soil. Noteworthy, the increase in Myxococcales occurred late in the incubation where especially Chitinophagaceae and Alphaproteobacteria decreased. Myxococcales are ‘micropredators’ and attack and lyse other bacteria which might explain the increased dominance of this group on the expense of other bacterial groups (Reichenbach, 1999).

The increase in 16S rRNA gene copy numbers after ash amendment (up to 12 t ha^−1^ and 90 t ha^−1^ for the agricultural and forest soil, respectively) is consistent with other reports of increasing bacterial numbers after wood ash application (Bååth & Arnebrant, 1994; Fritze *et al.*, 2000; Perkiömäki & Fritze, 2002; Bang-Andreasen *et al.*, 2017; Vestergård *et al.*, 2018). The large increase in the forest soil is further consistent with the increased pH as most bacteria thrive better at pH around 7 (Rousk *et al.*, 2009). Prokaryotic growth as well as a change towards a more copiotrophic community with higher average 16S rRNA gene number per genome is likely causing the 16S rRNA gene copy increase (Klappenbach *et al.*, 2000; Roller *et al.*, 2016).

The 90 t ha^−1^ ash amendment to the forest soil caused immediate dominance of Firmicutes and Gammaproteobacteria. Both groups are copiotrophs that thrives upon addition of easily degradable carbon and nitrogen to soil which probably partly explain their success upon ash application (Fierer *et al.*, 2007; Cleveland *et al.*, 2007; Nemergut *et al.*, 2010; Ramirez *et al.*, 2012; Fierer *et al.*, 2012). However, bacteria from these phyla are also known to be tolerant to heavy metals (Jacquiod *et al.*, 2017). Moreover, within Firmicutes the endospore-forming genus *Paenibacillus* dominated (de Hoon *et al.*, 2010), and we found increased transcription of genes involved in sporulation in these samples. Combined, these capabilities probably enable members of these groups to withstand the initial wood ash induced changes to the soil, including increased heavy metal concentrations, thereby allowing them to be initial utilizers of newly available labile resources. Reduced diversity at this ash dose further indicates that less organisms can cope with the ash induced changes to the soil system

### 4.2. Fungal responses to wood ash application

In both soil types, fungal response to ash amendment was slight compared to the prokaryotic response. Likewise, Cruz-Paredes et al. (2017), Högberg et al. (2007) and Rousk *et al.*, (2009, 2011) found bacteria to be more stimulated by nutrient addition and increases in pH than fungi. Similarly effects of ash amendment have been reported by Noyce *et al.* (2016) and Mahmood *et al.* (2003). The 90 t ha^−1^ amendment in the forest soil caused increased ITS gene copy numbers and a fungal community shift with increased dominance of *Mortierella, Peziza* and Hypocreales. These fungi are opportunistic saprotrophs with high growth rates and can exploit readily availible nutrients before other fungi arrive (Carlile *et al.*, 2001; Tedersoo *et al.*, 2006; Druzhinina *et al.*, 2012). Further, some *Peziza* spp. are early post-fire colonizers adapted to ash conditions (Egger, 1986; Rincón *et al.*, 2014). The increase in these groups further supports that copiotrophic-like lifestyles are favoured by wood ash application.

### 4.3. Micro-eukaryote responses to wood ash application

The micro-eukaryotes also responded to wood ash application in the forest soil, probably because the stimulation of copiotrophic bacteria and fungi provided more food for nematodes and protozoa (Rønn *et al.*, 2012). Ciliates (*Colpoda*), amoebae (Tubulinea) and small heterotrophic flagellates (Heteromitidae and Silicofilosea) increased with more pronounced responses at the later incubation times. Protozoa generally have longer generation times than prokaryotes, and thus need longer time to increase in population size. Further, they cannot start growth before a reasonable bacterial population has been formed (Fenchel, 1987; Ekelund *et al.*, 2002). The protozoan increase may explain the small decrease in prokaryotic 16S rRNA gene copies at day 100, where we observed the largest fraction of protozoa. The positively responding protozoa were likely primarily bacterivorous (Ekelund & Rønn, 1994; Ekelund, 1998), consistent with the decreasing relative fraction of bacterial rRNA sequences and the increasing relative fraction of fungal and protozoan rRNA sequences in the later incubation times after the application of 12 and 90 t ha^−1^ ash. Thus, preferential protozoan grazing on bacteria can explain the relative larger rRNA-fraction of fungi and protozoa at day 100. We found no significant effect of ash-amendment on micro-eukaryotes in the agricultural soil, which is consistent with the relative minor effects on prokaryotes and fungi in this soil.

### 4.5. Stress responses at high wood ash amendments

We recorded increased transcription of stress-response genes at the 90 t ha^−1^ amendments, which supports that this high dose exerts harmful effects on many members of the micro-biome. For example, chaperones ensure correct folding of proteins and are involved in cellular coping with stress-induced denaturation of proteins (Feder & Hofmann, 1999) and the increase in transcription level of these probably is a stress response. Also, transmembrane transporter proteins balance osmotic pressure of cells, regulate cytosolic pH and can export toxins such as metals from the cell (Alberts *et al.*, 2002; Ma *et al.*, 2009; Wilkens, 2015). Increased activity of transmembrane transporters is probably a response to wood ash induced osmotic changes to the soil system, increased pH, metal concentration and other toxic compounds.

### 4.6. The changes in the microbial communities are linked to physicochemical soil parameters

We found that ash-amendment strongly increased soil pH, which is a strong driver of microbial community composition and functioning (Fierer & Jackson, 2006; Rousk *et al.*, 2010) also after wood ash application (Frostegård *et al.*, 1993; Zimmermann & Frey, 2002; Högberg *et al.*, 2007; Peltoniemi *et al.*, 2016; Bang-Andreasen *et al.*, 2017). DOC and phosphate concomitantly increased. Several factors may contribute to this: (I) pH dependent changes in solubility (Evans *et al.*, 2012; Maresca *et al.*, 2017), (II) release from dead organisms incapable of coping with the wood ash or wood ash induced changes to the soil system, (III) increased mineralization rates after wood ash application (Bååth & Arnebrant, 1994; Vestergård *et al.*, 2018) and (IV) the phosphorous in the bio-ash (Pitman, 2006; Maresca *et al.*, 2017).

Since pH, conductivity, DOC and phosphate all correlated positively to wood ash concentrations it is difficult to disentangle the direct effect of these components as they might all be covariates of the wood ash amendments. pH-changes induce a cascade of effects in soil parameters and therefore affect mineral nutrient availability, salinity, metal solubility and organic C (Lauber *et al.*, 2009). Many of the wood ash induced changes were likely caused directly or indirectly by the pH increase, which is probably the major reason that pH is an essential driver of taxonomic and functional soil characteristics (Lauber *et al.*, 2009; Rousk *et al.*, 2010; Fierer, 2017; Vestergård *et al.*, 2018).

Wood ash contains virtually no nitrogen, hence measurable effects on soil nitrate and ammonium are probably caused by pH effects on microbial N mineralization (Vestergård *et al.*, 2018) and ion solubility (Pitman, 2006). Changes in nitrate and ammonium were significant as explanatory variables on the observed rRNA and mRNA dissimilarity profiles of the forest soil but not in the agricultural soil. Forest soil is generally more N limited than agricultural soil, where N is kept at a high level through fertilization.

## 4.7. Conclusions

We used detailed total RNA-Sequencing to demonstrate drastic taxonomic and functional changes in the active prokaryotic and eukaryotic micro-biomes of agricultural and forest soil after wood ash amendment. Our analyses suggested that increase in pH, electrical conductivity, dissolved organic carbon and phosphate were the main drivers of the observed changes. Wood ash amendment of 3 and 12 t ha^−1^ resulted in increased prokaryotic abundance and dominance of copiotrophic groups and elevated expression of genes involved in metabolism and cell growth. Amendment of 90 t ha^−1^ caused collapse of the micro-biome in the agricultural soil, while in the forest the copiotrophic micro-biome, also including fast-growing saprotrophic fungi, was further stimulated. However, diversity was reduced, and expression of stress response genes increased. Bacterivorous protozoan groups increased as a response to enhanced bacterial growth, which supports that the protozoa have a pivotal role in controlling bacterial abundance in soil following wood ash application. Overall, prokaryotic community and quantity responded more pronounced to wood ash amendment than fungi in both forest and agricultural soil.

## Supporting information

Supplementary Datasheet 1

Supplementary Datasheet 2

Supplementary Datasheet 3

Supplementary Datasheet 4

Supplementary Datasheet 5

## Acknowledgment

We thank Pia Bach Jakobsen for laboratory assistance.

## Figure Legends

**Supplementary Figure 1:**
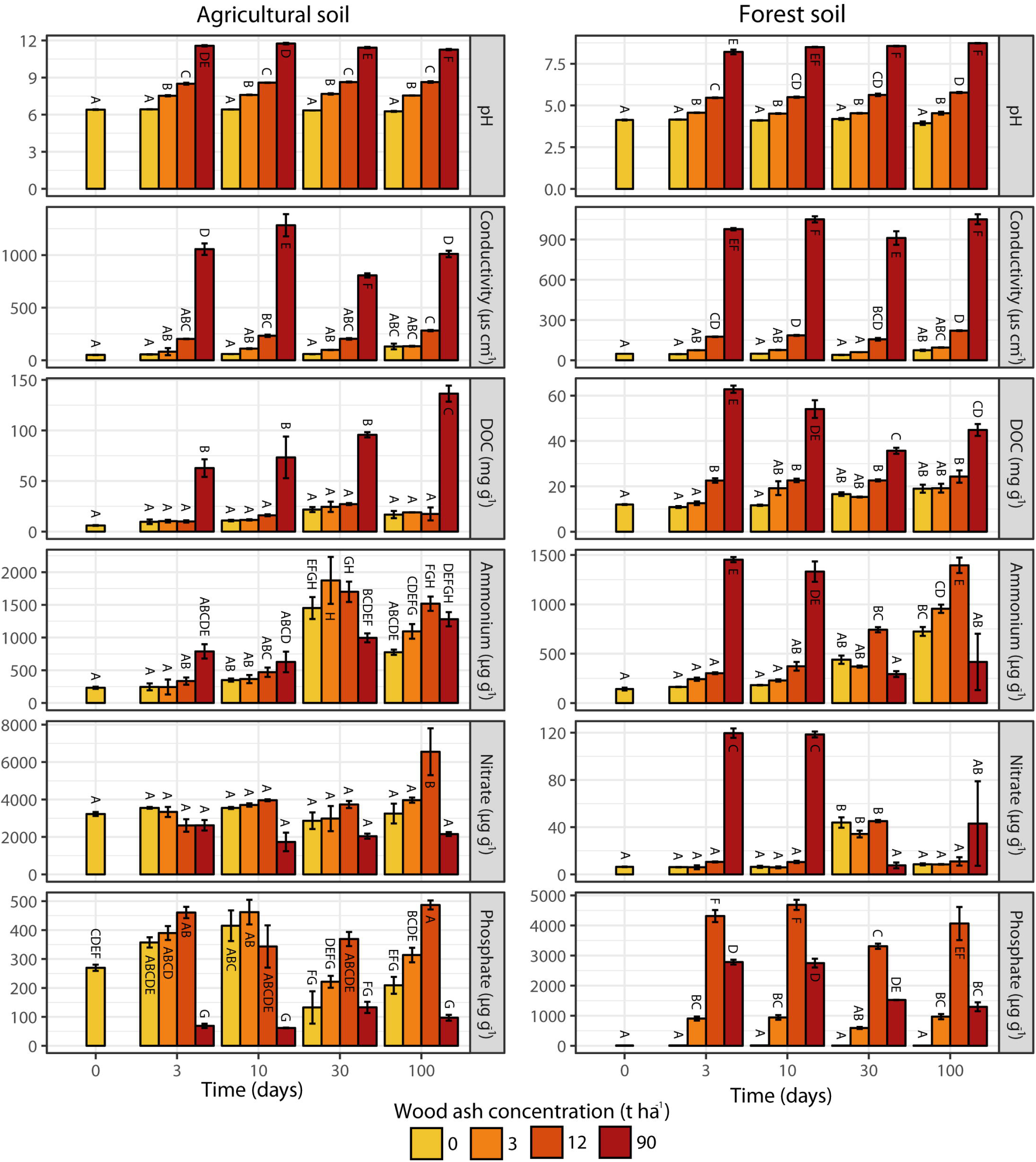
Metadata results across soil types, wood ash concentrations and incubation time. Different letters denote significant (p < 0.05) difference between samples within the same plot (Tukey post-hoc pairwise comparisons). Bars represents averages of triplicates with SEM (n = 3). Bars without errorbars represents values of 1 replicate. Note different range of y-axis values between the two soils for the same metadata category.

**Supplementary Figure 2:**
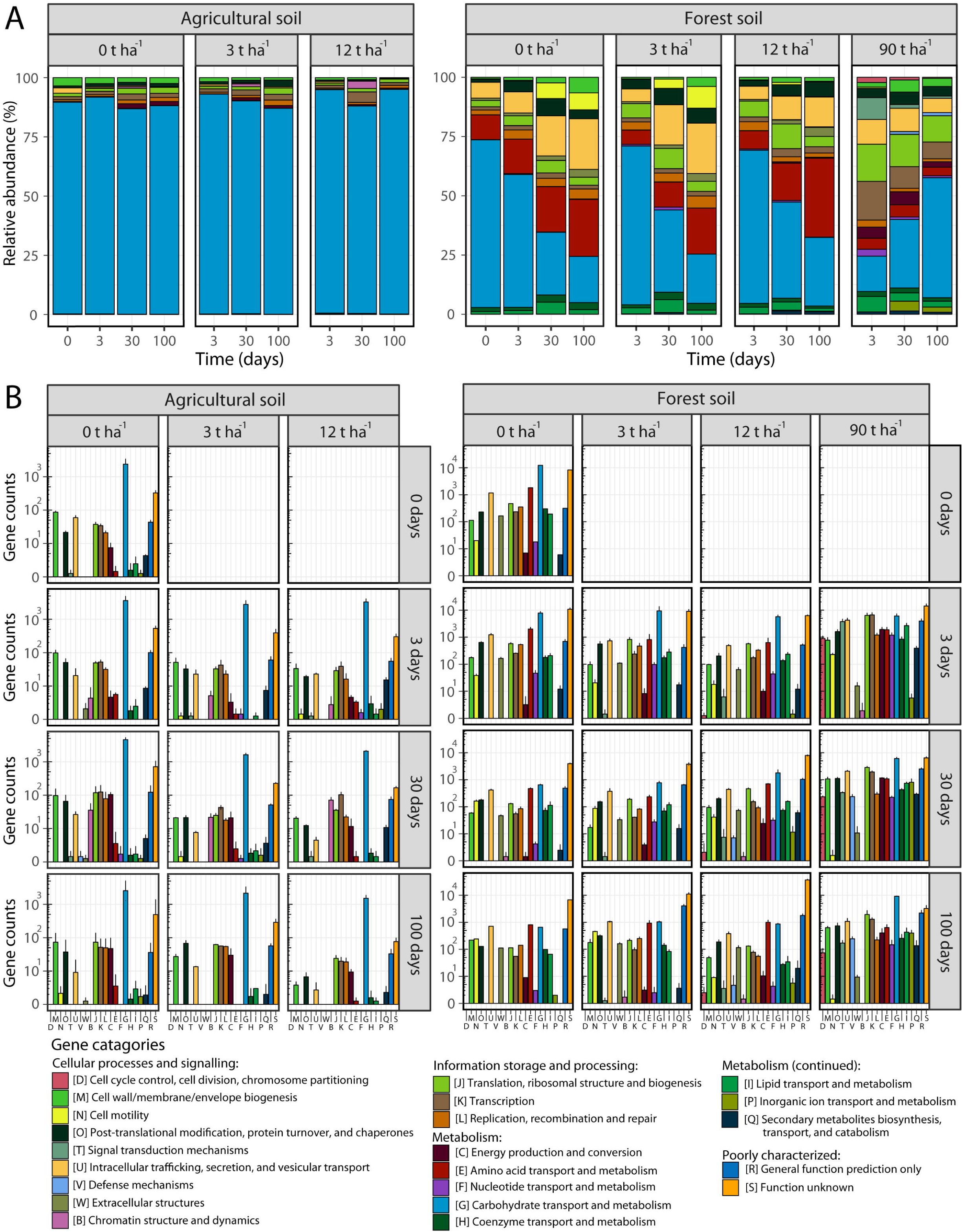
Functional gene compositions in (A) relative abundance and (B) absolute abundance (note log10 y-axis). “Poorly characterized” genes are excluded from the relative abundance plots to increase resolution of genes with known function. Bars are averages of triplicates with SEM as errorbars (excluding agricultural soil 3 t ha^−1^, 100 days (n=2) and forest soil 0t ha after 0 (n=1) and 100 days (n=1)).

**Supplementary Figure 3:**
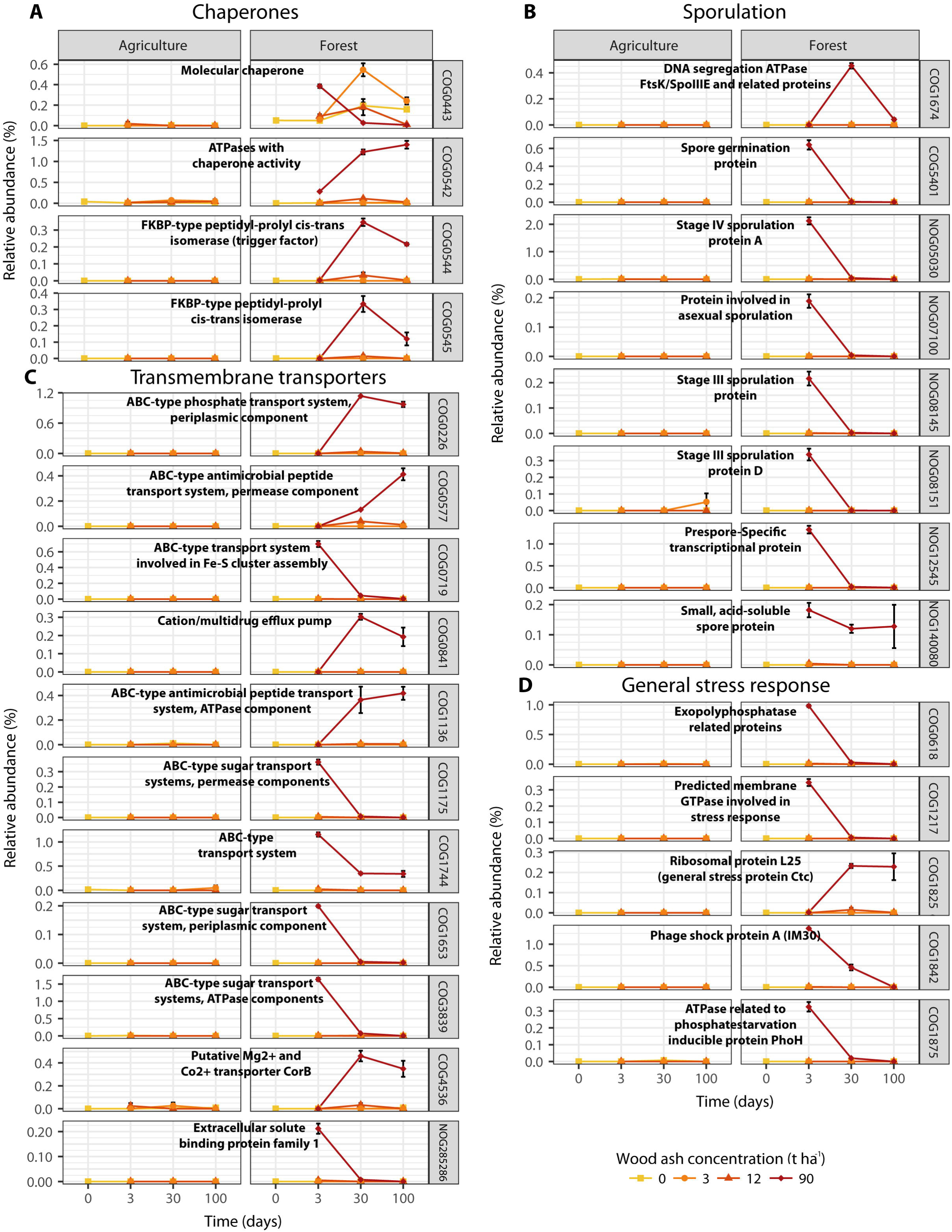
Functional genes involved in (A) chaperones, (B) sporulation, (C) Transmembrane transporters and (D) general stress response. The presented functional genes (with unique COG IDs) are all differentially expressed after wood ash amendment and are presented as relative abundance of total mRNA profile. Symbols are averages of triplicates with SEM as errorbars (excluding agricultural soil 3 t ha^−1^, 100 days (n=2) and forest soil 0t ha after 0 (n=1) and 100 days (n=1)).

## References

Alberts B, Johnson A, Lewis J, Raff M, Roberts K, Walter P. (2002). Molecular Biology of the Cell. 4th ed. Garland Science: New York.

Aronsson KA, Ekelund NGA. (2004). Biological effects of wood ash application to forest and aquatic ecosystems. J Environ Qual 33:1595–1605.

Augusto L, Bakker MR, Meredieu C. (2008). Wood ash applications to temperate forest ecosystems—potential benefits and drawbacks. Plant Soil 306:181–198.

Bang-Andreasen T, Nielsen JT, Voriskova J, Heise J, Rønn R, Kjøller R, et al. (2017). Wood ash induced pH changes strongly affect soil bacterial numbers and community composition. Front Microbiol 8:1400.

Bergmann GT, Bates ST, Eilers KG, Lauber CL, Caporaso JG, Walters WA, et al. (2011). The under-recognized dominance of Verrucomicrobia in soil bacterial communities. Soil Biol Biochem 43:1450–1455.

Blagodatskaya E, Kuzyakov Y. (2013). Active microorganisms in soil: Critical review of estimation criteria and approaches. Soil Biology and Biochemistry 67:192–211.

Blattner FR, Plunkett G, Bloch CA, Perna NT, Burland V, Riley M, et al. (1997). The complete genome sequence of Escherichia coli K-12. Science 277:1453–1462.

Bååth E, Arnebrant K. (1994). Growth rate and response of bacterial communities to pH in limed and ash treated forest soils. Soil Biology and Biochemistry 26:995–1001.

Carlile MJ, Watkinson SC, Gooday GW. (2001). The Fungi. 2nd ed. Academic Press: San Diego, Calif.

Carvalhais LC, Dennis PG, Tyson GW, Schenk PM. (2012). Application of metatranscriptomics to soil environments. J Microbiol Methods 91:246–251.

Cederlund H, Wessén E, Enwall K, Jones CM, Juhanson J, Pell M, et al. (2014). Soil carbon quality and nitrogen fertilization structure bacterial communities with predictable responses of major bacterial phyla. Applied Soil Ecology 84:62–68.

Cleveland CC, Nemergut DR, Schmidt SK, Townsend AR. (2007). Increases in soil respiration following labile carbon additions linked to rapid shifts in soil microbial community composition. Biogeochemistry 82:229–240.

Cookson WR, Osman M, Marschner P, Abaye DA, Clark I, Murphy DV, et al. (2007). Controls on soil nitrogen cycling and microbial community composition across land use and incubation temperature. Soil Biology and Biochemistry 39:744–756.

Cruz-Paredes C, Wallander H, Kjøller R, Rousk J. (2017). Using community trait-distributions to assign microbial responses to pH changes and Cd in forest soils treated with wood ash. Soil Biology and Biochemistry 112:153–164.

Demeyer A, Voundi Nkana J., Verloo M. (2001). Characteristics of wood ash and influence on soil properties and nutrient uptake: an overview. Bioresour Technol 77:287–295.

Druzhinina IS, Shelest E, Kubicek CP. (2012). Novel traits of Trichoderma predicted through the analysis of its secretome. FEMS Microbiol Lett 337:1–9.

Egger KN. (1986). Substrate Hydrolysis Patterns of Post-Fire Ascomycetes (Pezizales). Mycologia 78:771.

Ekelund F. (1998). Enumeration and abundance of mycophagous protozoa in soil, with special emphasis on heterotrophic flagellates. Soil Biology and Biochemistry 30:1343–1347.

Ekelund F, Frederiksen HB, Rønn R. (2002). Population dynamics of active and total ciliate populations in arable soil amended with wheat. Appl Environ Microbiol 68:1096–1101.

Ekelund F, Rønn R. (1994). Notes on protozoa in agricultural soil with emphasis on heterotrophic flagellates and naked amoebae and their ecology. FEMS Microbiol Rev 15:321–353.

Epelde L, Lanzén A, Blanco F, Urich T, Garbisu C. (2015). Adaptation of soil microbial community structure and function to chronic metal contamination at an abandoned Pb-Zn mine. FEMS Microbiol Ecol 91:1–11.

Evans CD, Jones TG, Burden A, Ostle N, Zieliński P, Cooper MDA, et al. (2012). Acidity controls on dissolved organic carbon mobility in organic soils. Glob Change Biol 18:3317–3331.

Feder ME, Hofmann GE. (1999). Heat-shock proteins, molecular chaperones, and the stress response: evolutionary and ecological physiology. Annu Rev Physiol 61:243–282.

Fenchel T. (1987). Ecology of Protozoa. 1st ed. Springer: Berlin, Heidelberg doi:10.1007/978-3-662-25981-8.

Fierer N. (2017). Embracing the unknown: disentangling the complexities of the soil microbiome. Nat Rev Microbiol 15:579–590.

Fierer N, Bradford MA, Jackson RB. (2007). Toward an ecological classification of soil bacteria. Ecology 88:1354–1364.

Fierer N, Jackson RB. (2006). The diversity and biogeography of soil bacterial communities. Proc Natl Acad Sci U S A 103:626–631.

Fierer N, Lauber CL, Ramirez KS, Zaneveld J, Bradford MA, Knight R. (2012). Comparative metagenomic, phylogenetic and physiological analyses of soil microbial communities across nitrogen gradients. ISME J 6:1007–1017.

Fritze H, Perkiömäki J, Saarela U, Katainen R, Tikka P, Yrjälä K, et al. (2000). Effect of Cd-containing wood ash on the microflora of coniferous forest humus. FEMS Microbiol Ecol 32:43–51.

Frostegård A, Tunlid A, Bååth E. (1993). Phospholipid Fatty Acid composition, biomass, and activity of microbial communities from two soil types experimentally exposed to different heavy metals. Appl Environ Microbiol 59:3605–3617.

Frostegård Å., Bååth E, Tunlio A. (1993). Shifts in the structure of soil microbial communities in limed forests as revealed by phospholipid fatty acid analysis. Soil Biology and Biochemistry 25:723–730.

Geisen S, Tveit AT, Clark IM, Richter A, Svenning MM, Bonkowski M, et al. (2015). Metatranscriptomic census of active protists in soils. ISME J 9:2178–2190.

Grabherr MG, Haas BJ, Yassour M, Levin JZ, Thompson DA, Amit I, et al. (2011). Full-length transcriptome assembly from RNA-Seq data without a reference genome. Nat Biotechnol 29:644– 652.

Hanada S, Tamaki H, Nakamura K, Kamagata Y. (2014). Crenotalea thermophila gen. nov., sp. nov., a member of the family Chitinophagaceae isolated from a hot spring. Int J Syst Evol Microbiol 64:1359–1364.

Hansen CHF, Krych L, Nielsen DS, Vogensen FK, Hansen LH, Sørensen SJ, et al. (2012). Early life treatment with vancomycin propagates Akkermansia muciniphila and reduces diabetes incidence in the NOD mouse. Diabetologia 55:2285–2294.

Hansen M, Bang-Andreasen T, Sørensen H, Ingerslev M. (2017). Micro vertical changes in soil pH and base cations over time after application of wood ash on forest soil. Forest Ecology and Management 406:274–280.

De Hoon MJL, Eichenberger P, Vitkup D. (2010). Hierarchical evolution of the bacterial sporulation network. Curr Biol 20:R735–45.

Hultman J, Waldrop MP, Mackelprang R, David MM, McFarland J, Blazewicz SJ, et al. (2015). Multi-omics of permafrost, active layer and thermokarst bog soil microbiomes. Nature 521:208–212.

Huotari N, Tillman-Sutela E, Moilanen M, Laiho R. (2015). Recycling of ash – For the good of the environment? Forest Ecology and Management 348:226–240.

Högberg MN, Högberg P, Myrold DD. (2007). Is microbial community composition in boreal forest soils determined by pH, C-to-N ratio, the trees, or all three? Oecologia 150:590–601.

Ihrmark K, Bödeker ITM, Cruz-Martinez K, Friberg H, Kubartova A, Schenck J, et al. (2012). New primers to amplify the fungal ITS2 region--evaluation by 454-sequencing of artificial and natural communities. FEMS Microbiol Ecol 82:666–677.

Jacquiod S, Cyriaque V, Riber L, Al-soud WA, Gillan DC, Wattiez R, et al. (2017). Long-term industrial metal contamination unexpectedly shaped diversity and activity response of sediment microbiome. J Hazard Mater 344:299–307.

Jensen LJ, Julien P, Kuhn M, von Mering C, Muller J, Doerks T, et al. (2008). eggNOG: automated construction and annotation of orthologous groups of genes. Nucleic Acids Res 36:D250–4.

Karltun E, Saarsalmi A, Ingerslev M, Mandre M, Andersson S, Gaitnieks T, et al. (2008). Wood ash recycling – possibilities and risks. In:Sustainable use of forest biomass for energy, Röser, D, Asikainen, A, Raulund-Rasmussen, K, & Stupak, I (eds), Springer Netherlands: Dordrecht, pp. 79–108.

Kielak AM, Barreto CC, Kowalchuk GA, van Veen JA, Kuramae EE. (2016). The Ecology of Acidobacteria: Moving beyond Genes and Genomes. Front Microbiol 7:744.

Kim JM, Roh A-S, Choi S-C, Kim E-J, Choi M-T, Ahn B-K, et al. (2016). Soil pH and electrical conductivity are key edaphic factors shaping bacterial communities of greenhouse soils in Korea. J Microbiol 54:838–845.

Klappenbach JA, Dunbar JM, Schmidt TM. (2000). rRNA operon copy number reflects ecological strategies of bacteria. Appl Environ Microbiol 66:1328–1333.

Kopylova E, Noé L, Touzet H. (2012). SortMeRNA: fast and accurate filtering of ribosomal RNAs in metatranscriptomic data. Bioinformatics 28:3211–3217.

Kämpfer P, Young C-C, Sridhar KR, Arun AB, Lai WA, Shen FT, et al. (2006). Transfer of [Flexibacter] sancti, [Flexibacter] filiformis, [Flexibacter] japonensis and [Cytophaga] arvensicola to the genus Chitinophaga and description of Chitinophaga skermanii sp. nov. Int J Syst Evol Microbiol 56:2223–2228.

Lanzén A, Jørgensen SL, Huson DH, Gorfer M, Grindhaug SH, Jonassen I, et al. (2012). CREST--classification resources for environmental sequence tags. PLoS ONE 7:e49334.

Lauber CL, Hamady M, Knight R, Fierer N. (2009). Pyrosequencing-based assessment of soil pH as a predictor of soil bacterial community structure at the continental scale. Appl Environ Microbiol 75:5111–5120.

Li H, Durbin R. (2009). Fast and accurate short read alignment with Burrows-Wheeler transform. Bioinformatics 25:1754–1760.

Lladó S, Baldrian P. (2017). Community-level physiological profiling analyses show potential to identify the copiotrophic bacteria present in soil environments. PLoS ONE 12:e0171638.

Love MI, Huber W, Anders S. (2014). Moderated estimation of fold change and dispersion for RNA-seq data with DESeq2. Genome Biol 15:550–550.

Ma Z, Jacobsen FE, Giedroc DP. (2009). Metal Transporters and Metal Sensors: How Coordination Chemistry Controls Bacterial Metal Homeostasis. Chem Rev 109:4644–4681.

Mahmood S, Finlay RD, Fransson A-M, Wallander H. (2003). Effects of hardened wood ash on microbial activity, plant growth and nutrient uptake by ectomycorrhizal spruce seedlings. FEMS Microbiol Ecol 43:121–131.

Maresca A, Hyks J, Astrup TF. (2017). Recirculation of biomass ashes onto forest soils: ash composition, mineralogy and leaching properties. Waste Manag 70:127–138.

Martin M. (2011). Cutadapt removes adapter sequences from high-throughput sequencing reads. EMBnet.journal 17:10.

Miller CS, Baker BJ, Thomas BC, Singer SW, Banfield JF. (2011). EMIRGE: reconstruction of full-length ribosomal genes from microbial community short read sequencing data. Genome Biol 12:R44.

Nawrocki EP, Burge SW, Bateman A, Daub J, Eberhardt RY, Eddy SR, et al. (2015). Rfam 12.0: updates to the RNA families database. Nucleic Acids Res 43:D130–7.

Nemergut DR, Cleveland CC, Wieder WR, Washenberger CL, Townsend AR. (2010). Plot-scale manipulations of organic matter inputs to soils correlate with shifts in microbial community composition in a lowland tropical rain forest. Soil Biology and Biochemistry 42:2153–2160.

Noyce GL, Fulthorpe R, Gorgolewski A, Hazlett P, Tran H, Basiliko N. (2016). Soil microbial responses to wood ash addition and forest fire in managed Ontario forests. Applied Soil Ecology 107:368–380.

Ohno T, Susan Erich M. (1990). Effect of wood ash application on soil pH and soil test nutrient levels. Agriculture, Ecosystems & Environment 32:223–239.

Oksanen J, Kindt R, Legendre P, O’Hara B, Simpson GL, Solymos P, et al. (2008). vegan: Community Ecology Package. https://cran.r-project.org/web/packages/vegan/.

Peltoniemi K, Pyrhönen M, Laiho R, Moilanen M, Fritze H. (2016). Microbial communities after wood ash fertilization in a boreal drained peatland forest. European journal of soil biology 76:95– 102.

Perkiömäki J, Fritze H. (2002). Short and long-term effects of wood ash on the boreal forest humus microbial community. Soil Biology and Biochemistry 34:1343–1353.

Pitman RM. (2006). Wood ash use in forestry - a review of the environmental impacts. Forestry 79:563–588.

Qin J, Hovmand MF, Ekelund F, Rønn R, Christensen S, Groot GA de, et al. (2017). Wood ash application increases pH but does not harm the soil mesofauna. Environ Pollut 224:581–589.

R Core Team. (2015). R: A language and environment for statistical computing. http://www.R-project.org/.

Ramirez KS, Craine JM, Fierer N. (2012). Consistent effects of nitrogen amendments on soil microbial communities and processes across biomes. Glob Change Biol 18:1918–1927.

Reichenbach H. (1999). The ecology of the myxobacteria. Environ Microbiol 1:15–21.

Rice P, Longden I, Bleasby A. (2000). EMBOSS: the european molecular biology open software suite. Trends Genet 16:276–277.

Rincón A, Santamaría BP, Ocaña L, Verdú M. (2014). Structure and phylogenetic diversity of post-fire ectomycorrhizal communities of maritime pine. Mycorrhiza 24:131–141.

Roller BRK, Stoddard SF, Schmidt TM. (2016). Exploiting rRNA operon copy number to investigate bacterial reproductive strategies. Nature microbiology 1:16160.

Rousk J, Brookes PC, Bååth E. (2009). Contrasting soil pH effects on fungal and bacterial growth suggest functional redundancy in carbon mineralization. Appl Environ Microbiol 75:1589–1596.

Rousk J, Brookes PC, Bååth E. (2011). Fungal and bacterial growth responses to N fertilization and pH in the 150-year “Park Grass” UK grassland experiment. FEMS Microbiol Ecol 76:89–99.

Rousk J, Bååth E, Brookes PC, Lauber CL, Lozupone C, Caporaso JG, et al. (2010). Soil bacterial and fungal communities across a pH gradient in an arable soil. ISME J 4:1340–1351.

Rønn R, Vestergård M, Ekelund F. (2012). Interactions Between Bacteria, Protozoa and Nematodes in Soil. Acta Protozoologica 51:223–235.

Schostag M, Priemé A, Jacquiod R, Russel J, Ekelund F, Jacobsen CS. (2019). Bacterial and protozoan dynamics upon thawing and freezing of an active layer permafrost soil. ISME J doi:10.1038/s41396-019-0351-x.

Starke R, Kermer R, Ullmann-Zeunert L, Baldwin IT, Seifert J, Bastida F, et al. (2016). Bacteria dominate the short-term assimilation of plant-derived N in soil. Soil Biology and Biochemistry 96:30–38.

Tedersoo L, Hansen K, Perry BA, Kjøller R. (2006). Molecular and morphological diversity of pezizalean ectomycorrhiza. New Phytol 170:581–596.

Teng Y, Wang X, Li L, Li Z, Luo Y. (2015). Rhizobia and their bio-partners as novel drivers for functional remediation in contaminated soils. Front Plant Sci 6:32.

Urich T, Lanzén A, Qi J, Huson DH, Schleper C, Schuster SC. (2008). Simultaneous assessment of soil microbial community structure and function through analysis of the meta-transcriptome. PLoS ONE 3:e2527.

Vance ED. (1996). Land Application of Wood-Fired and Combination Boiler Ashes: An Overview. Journal of Environment Quality 25:937.

Varet H, Brillet-Guéguen L, Coppée J-Y, Dillies M-A. (2016). SARTools: A DESeq2- and EdgeR- Based R Pipeline for Comprehensive Differential Analysis of RNA-Seq Data. PLoS ONE 11:e0157022.

Vaser R, Pavlović D, Šikić M. (2016). SWORD-a highly efficient protein database search. Bioinformatics 32:i680–i684.

Vestergård M, Bang-Andreasen T, Buss SM, Cruz-Paredes C, Bentzon-Tilia S, Ekelund F, et al. (2018). The relative importance of the bacterial pathway and soil inorganic nitrogen increase across an extreme wood-ash application gradient. Glob Change Biol Bioenergy. doi:10.1111/gcbb.12494.

Wall DH, Bardgett RD, Behan-Pelletier V, Herrick JE, Jones TH, Ritz K, et al. (2012). Soil ecology and ecosystem services. 1st ed. Oxford University Press: USA.

White T, Bruns R, Lee S, Taylor J. (1990). Amplification and direct sequencing of fungal ribosomalrna genes for phylogenetics. In:PCR protocols - a guide to methods and applications, Innis, M, Gelfand, D, Sninsky, J, &White, T (eds), Academic Press: New York, US, pp. 315–322.

Wilke A, Harrison T, Wilkening J, Field D, Glass EM, Kyrpides N, et al. (2012). The M5nr: a novel non-redundant database containing protein sequences and annotations from multiple sources and associated tools. BMC Bioinformatics 13:141.

Wilkens S. (2015). Structure and mechanism of ABC transporters. F1000Prime Rep 7:14.

Zimmermann S, Frey B. (2002). Soil respiration and microbial properties in an acid forest soil: effects of wood ash. Soil Biology and Biochemistry 34:1727–1737.

